# A natural glycolipid exposes the outer membrane interface as a tunable regulator of bacterial surface attachment

**DOI:** 10.64898/2026.08.20.745940

**Authors:** Han-Yi Huang, Elsa N. Astorga-Simon, Christopher Adamson, Yuan Qiao, Paula P. Navarro, Alexandre Persat

## Abstract

Surface attachment initiates bacterial colonization and biofilm formation, yet remains difficult to target owing to redundant and species-specific mechanisms. Here we identify dalberoside, a natural glycolipid that inhibits adhesion by non-disruptive remodeling of the Gram-negative outer membrane interface. Dalberoside inhibited *Pseudomonas aeruginosa* and *Acinetobacter baumannii* attachment and biofilm formation without measurable bactericidal activity or membrane permeabilization. It reduced *P. aeruginosa* retention on epithelial cells, delayed *P. aeruginosa* cytotoxicity and rapidly detached surface-associated *A. baumannii*. High-resolution microscopy of a fluorescent analogue and *in situ* cryo-electron tomography support a model in which dalberoside associates with the bacterial outer membrane, remodeling its outer leaflet. Biophysical measurements further indicated altered local interfacial properties. Thus, dalberoside reveals the outer membrane interface as a chemically addressable regulator of bacterial surface attachment.

## Introduction

Surface association is an early commitment step in bacterial colonization, enabling cells to persist on abiotic materials, establish biofilms and engage host tissues during infection.^1^ Surface-associated populations are more resilient to clearance and become more tolerant to antimicrobial treatment, making the earliest stages of attachment an attractive point of intervention.^2^ However, approaches that prevent bacterial surface association without inhibiting growth remain limited. Such non-bactericidal interventions represent a form of anti-virulence strategy, aiming to reduce colonization and persistence rather than directly eliminate bacterial cells.^3,4^

Anti-attachment strategies target defined adhesins, adhesin–receptor interactions or intracellular signaling pathways that regulate biofilm-associated behaviors.^5–13^ These approaches can be highly effective when attachment depends on a dominant molecular interaction. By contrast, opportunistic Gram-negative pathogens such as *Pseudomonas aeruginosa* and *Acinetobacter baumannii* adhere to diverse biotic and abiotic surfaces through multiple, partially redundant mechanisms, promoting persistence and contributing to treatment failure in biofilm-associated infections.^1,14–16^ This breadth complicates the development of broadly active anti-attachment agents and raises the possibility that shared physical properties of the bacterial cell surface, rather than individual adhesins, could provide an alternative point of control.^17–19^

The bacterial cell surface forms the interface through which cells encounter materials, host tissues, and neighboring cells.^20^ Its molecular composition and physicochemical properties govern the short-range interactions required for stable surface association.^20^ In Gram-negative bacteria, the outer membrane mediates these environmental interactions while functioning as an essential permeability barrier.^21^ Because perturbation of the cell surface often compromises barrier function, most outer membrane-targeting agents, such as polymyxins, are bactericidal.^22,23^ Whether cell-surface properties can instead be modulated to inhibit attachment while preserving envelope integrity therefore remains unclear.^3^ Natural products provide a structurally diverse source of molecules for probing such mechanisms, including amphiphilic metabolites that could interact with cellular interfaces and may enable alternative modes of surface modulation.^24^

Here, we developed an image-based phenotypic screen to identify natural products that reduce surface attachment of *P. aeruginosa* without prior knowledge of a molecular target. We identified dalberoside (DBO), a plant-derived glycolipid that broadly inhibits attachment of Gram-negative bacteria without impairing their viability. We then used DBO as a chemical probe to ask whether adhesion can be controlled by modifying the bacterial cell surface while preserving the essential barrier function of the outer membrane. By following its effects from initial attachment to biofilms and host-cell interactions, and combining chemical dissection with cellular imaging, cryo-electron tomography and biophysical measurements, we connect this phenotype to non-disruptive remodeling of the outer membrane. Our findings expose this interface as an underexplored point of control for surface-associated bacterial behaviors.

## Results

### Dalberoside inhibits *P. aeruginosa* and *A. baumannii* surface attachment

To identify compounds that inhibit bacterial surface attachment, we developed an image-based screen targeting *P. aeruginosa* strain PAO1 as a model organism that relies on strong surface attachment for colonization and infection (Fig. 1a, Extended Data Fig. 1). Wild-type (WT) cells were allowed to attach for 30 min in the presence of each compound of a natural product library, and then washed to quantify the stably attached population. We compared attachment of WT with a Δ*psl* mutant, which lacks an extracellular-polysaccharide (EPS) matrix component and exhibits much lower attachment than WT. In parallel, we screened the same library in the Δ*psl* mutant background, allowing us to identify compounds that were EPS-independent.

**Fig. 1.**
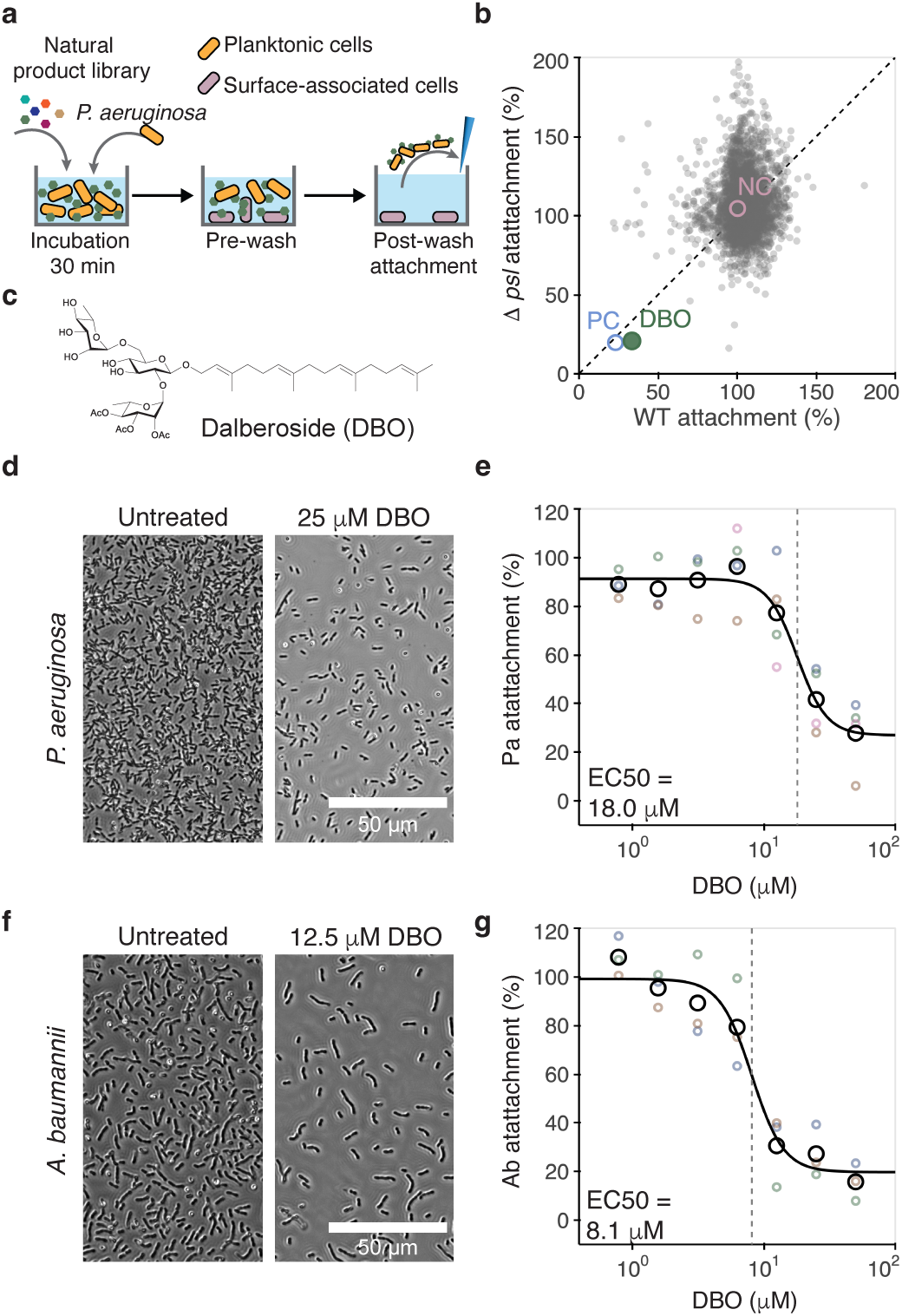
Identification of dalberoside (DBO) as an inhibitor of *Pseudomonas aeruginosa* and *Acinetobacter baumannii* surface attachment. **a,** Image-based phenotypic anti-attachment screening workflow. *P. aeruginosa* wild-type (WT) cells were incubated with library compounds for 30 min and washed to remove cells not stably associated with the surface, whereas Δ*psl* cells were imaged without washing. **b,** Screen of 2,627 natural products at 30 µM. Gray dots represent individual compounds. WT post-wash attachment (x axis) and Δ*psl* pre-wash attachment (y axis) were normalized to DMSO controls. NC denotes DMSO; PC denotes Δ*psl* in the post-wash assay and Δ*psl*Δ*pilA* in the pre-wash assay. Controls were selected by Z′-factor benchmarking (Extended Data Fig. 1). **c,** Chemical structure of DBO. **d,f,** Representative post-wash images of *P. aeruginosa* (**d**) and *A. baumannii* (**f**) after a 30-min treatment with DMSO or DBO. **e,g,** Concentration–response analysis in *P. aeruginosa* (**e**) and *A. baumannii* (**g**). Attachment was quantified as total segmented cell area per image and normalized to the mean DMSO control within each biological replicate. Colored open circles indicate biological-replicate means, black open circles indicate the mean across replicates and black curves indicate four-parameter log-logistic fits to biological-replicate means. Gray dashed lines indicate fitted EC50 values. Scale bars, 50 µm. For **e** and **g**, *n* = 4 and 3 independent biological replicates, respectively.

One compound reduced bacterial surface attachment by more than 50% in both screening conditions (Fig. 1b). We named this natural glycolipid dalberoside after *Dalbergia oliveri*, the plant from which it was isolated. Dalberoside is composed of a β-glucose core, an O-acetylated rhamnose, a second rhamnose, and a geranylgeranyl lipid tail (Fig. 1c). Dalberoside reduced *P. aeruginosa* attachment in a concentration-dependent manner, with an EC50 of 18.0 µM (Fig. 1d, e). It was also efficient against *A. baumannii* attachment, with an EC50 of 8.1 µM (Fig. 1f, g). A high-purity synthetic sample of dalberoside, prepared through the total synthesis reported in a companion paper, displayed greater anti-attachment potency with EC50 values of 9.1 µM for *P. aeruginosa* and 2.8 µM for *A. baumannii*.^25^ Together, these results establish dalberoside as a concentration-dependent inhibitor of surface attachment in both species.

### Dalberoside inhibits attachment without bactericidal activity

We next asked whether reduced bacterial viability contributed to the decrease in surface attachment. Dalberoside did not completely inhibit growth of either strain over the accessible concentration range, and therefore minimum inhibitory concentration (MIC) could be determined (Fig. 2a–d, Extended Data Fig. 2). At concentrations above those required to inhibit attachment (>25 µM), area under the curve analysis showed dalberoside modestly reduced growth, with reductions remaining under 30% for both strains. Consistent with the absence of bactericidal activity, treatment with 25 µM dalberoside, a concentration used in subsequent experiments, did not reduce CFU counts in either *P. aeruginosa* or *A. baumannii* (Fig. 2e).

**Fig. 2.**
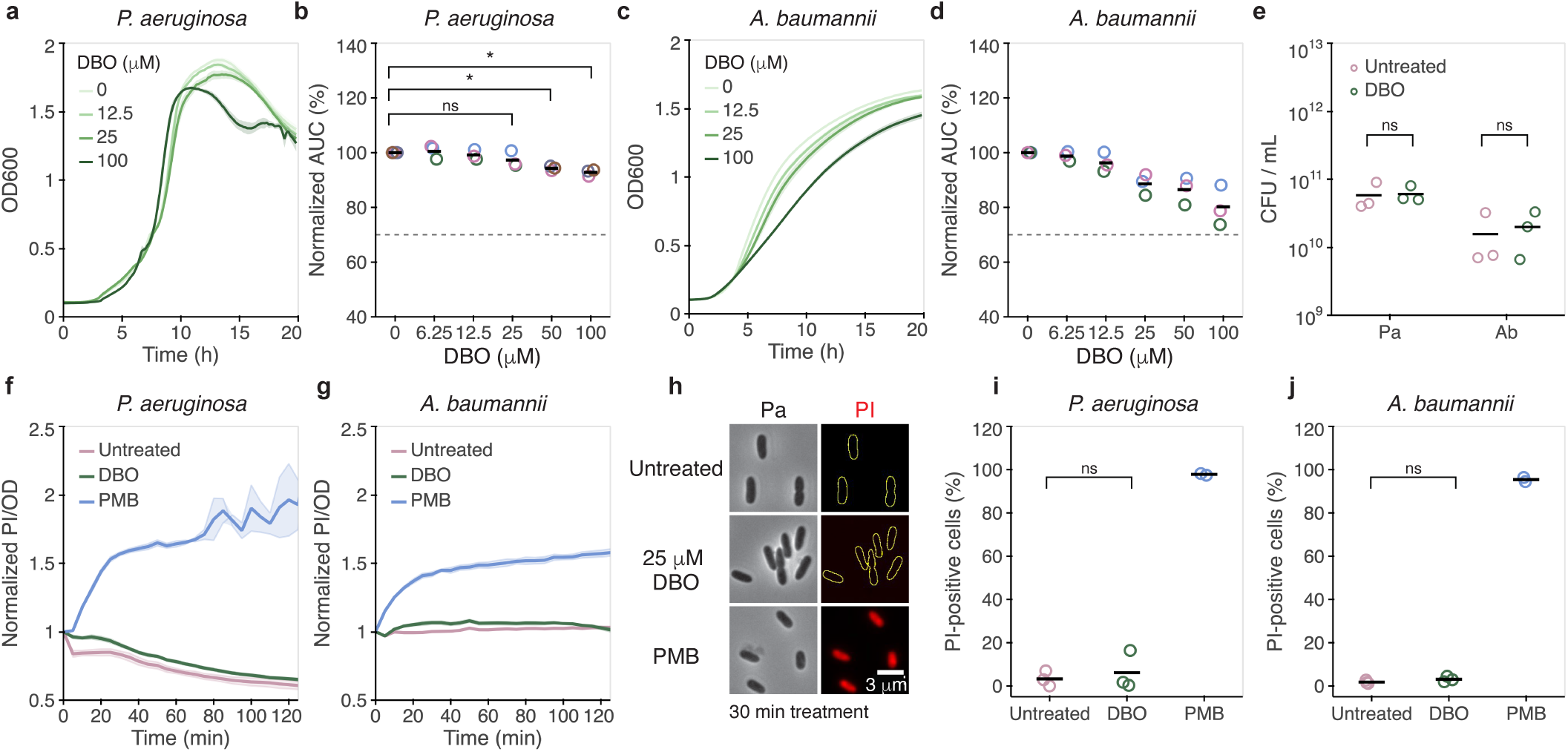
Dalberoside (DBO) does not impair bacterial growth, viability or membrane integrity at anti-attachment concentrations. **a–d,** Representative growth curves (**a,c**) and normalized area-under-the-curve (AUC) analyses (**b,d**) of *Pseudomonas aeruginosa* (**a,b**) and *Acinetobacter baumannii* (**c,d**) treated with the indicated DBO concentrations in Mueller–Hinton broth for 20 h. AUC values were calculated from the OD₆₀₀ growth curves and normalized to the mean DMSO control within each biological replicate. The gray dashed lines indicate the 70% normalized AUC. **e,** CFU enumeration after treatment with DMSO or 25 µM DBO for 24 h in LB at 37 °C with shaking. **f,g,** Representative time courses of PI fluorescence normalized to OD₆₀₀ and the initial value within each well for *P. aeruginosa* (**f**) and *A. baumannii* (**g**) treated with DMSO, 25 µM DBO or polymyxin B (PMB; 8 µg/ml) for 2 h. Lines and shaded regions indicate the mean ± SEM of technical replicate wells. **h,** Representative PI images of *P. aeruginosa* treated with DMSO, 25 µM DBO or PMB (100 µg/ml) for 30 min. Scale bars, 3 µm. **i,j,** PI-positive fractions in *P. aeruginosa* (**i**) and *A. baumannii* (**j**) after the treatments described in **h**. In **b,d,e,i,j**, colored open circles represent independent biological replicates and black horizontal lines indicate the mean; *n* = 3. Panels **a,c,f–h** show representative data from one of three independent biological replicates. Statistical analyses are described in Methods. *P* < 0.05, *; *P* < 0.01, **; *P* < 0.001, ***; *P* < 0.0001, ****; ns, not significant.

To determine whether the reduction in attachment was associated with sublethal membrane permeabilization, we next assessed propidium iodide (PI) uptake under conditions that inhibited attachment without reducing bacterial viability. Bulk PI fluorescence remained indistinguishable between the dalberoside-treated and vehicle-control groups throughout the 2 h monitoring period (Fig. 2f, g). Similarly, treating *P. aeruginosa* with dalberoside for 30 min did not measurably increase the proportion of PI-positive cells (Fig. 2h–j). Thus, dalberoside inhibits surface attachment without reducing bacterial viability or compromising membrane permeability.

### Dalberoside anti-biofilm activity

To form biofilms, bacteria must first robustly attach to surfaces. We therefore asked whether dalberoside inhibits biofilm formation and disrupts preformed biofilms. We used crystal violet staining as a quantitative readout of biofilm biomass formed under static conditions in the presence of the compound (Fig. 3a).^26^ Dalberoside reduced biofilm biomass in both species in a concentration-dependent manner (Fig. 3b, c). At 25 µM, a concentration at which dalberoside significantly decreased biofilm biomass, the proportion of viable cells retained on the surface decreased in both species, without reducing the total recoverable CFU within each well (Fig. 3b–d, Extended Data Fig. 3). These results suggest that dalberoside limits biofilm accumulation by reducing surface retention rather than bacterial growth.

**Fig. 3.**
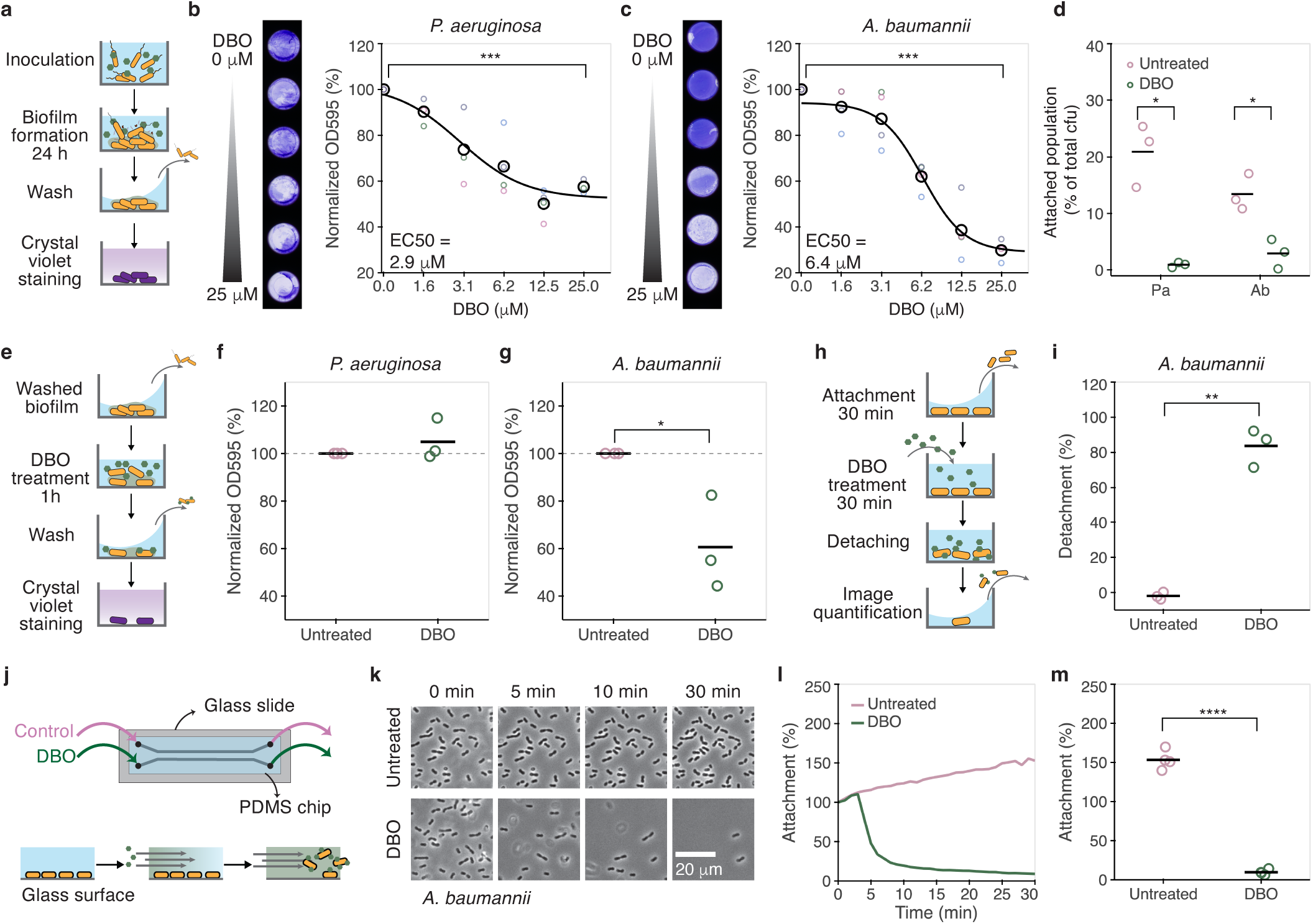
Dalberoside (DBO) inhibits biofilm formation and promotes detachment of surface-associated *A. baumannii*. **a,** Static biofilm-formation assay workflow. Bacteria were incubated with DMSO or DBO for 24 h before crystal violet staining. **b,c,** Representative stained biofilms and concentration–response analyses of *Pseudomonas aeruginosa* (**b**) and *A. baumannii* (**c**). OD₅₉₅ values were normalized to the DMSO control within each biological replicate. Colored open circles indicate biological-replicate means, black open circles indicate the mean across replicates, black curves indicate four-parameter log-logistic fits and fitted EC50 values are indicated. **d,** Surface-associated CFUs after 24 h incubation with DMSO or 25 µM DBO, expressed as a percentage of total recoverable CFUs per well. **e,** Preformed-biofilm reduction assay workflow. Biofilms were established for 23 h and treated for 1 h. **f,g,** Preformed-biofilm biomass after treatment with DMSO or 25 µM DBO in *P. aeruginosa* (**f**) and *A. baumannii* (**g**), normalized to DMSO. **h,** Plate-based detachment assay workflow. **i,** DBO-induced detachment of *A. baumannii*, calculated from pre- and post-treatment segmented cell areas from the same well. **j,** Microfluidic detachment assay schematic. **k,l,** Representative images (**k**) and time course (**l**) of *A. baumannii* during perfusion with DMSO or 25 µM DBO. Values were normalized to the initial time point. Scale bars, 20 µm. **m,** Surface-associated bacteria remaining after 30 min of perfusion. Points indicate independent biological replicates and black horizontal lines indicate the mean. For **b,c,m**, *n* = 4; for **d,f,g,i**, *n* = 3 independent biological replicates. Panels **k,l** show one representative experiment from four biological replicates. Statistical analyses are described in Methods. *P* < 0.05, *; *P* < 0.01, **; *P* < 0.001, ***; *P* < 0.0001, ****; ns, not significant.

We next examined whether dalberoside could disrupt already formed biofilms. We treated biofilms for 1 h before crystal violet quantification (Fig. 3e). Dalberoside at 25 µM reduced preformed *A. baumannii* biofilm biomass by approximately 40%. However, it caused no detectable change in *P. aeruginosa* crystal violet signal (Fig. 3f, g), revealing a species-dependent effect on already-formed biofilms. To determine whether this difference reflected direct loss of cell–surface retention, we next tested the detachment of single surface-associated bacteria.

To test its effect on detachment, we incubated pre-attached cells with dalberoside for 30 min before washing and quantification (Fig. 3h). Dalberoside detached more than 80% of surface-associated *A. baumannii* cells while we could not measure significant detachment of *P. aeruginosa* cells (Fig. 3i; Extended Data Fig. 4a, b). Because the plate-based assay provided only an endpoint measurement, we next used a microfluidic flow assay to resolve the kinetics of dalberoside-induced detachment under steady, controlled shear forces (Fig. 3j). Time-lapse imaging under continuous flow of dalberoside showed that *A. baumannii* rapidly detached in minutes (Fig. 3k, l). Surface coverage fell below 10% of its initial level within 30 min (Fig. 3m). In contrast, we could not measure any significant detachment of *P. aeruginosa* during 60 min of treatment (Extended Data Fig. 4c, d). Together, these results show that dalberoside inhibits biofilm formation in both species but is more potent in *A. baumannii* where it promotes the detachment of established surface-associated cells and biofilm.

### Dalberoside reduces *P. aeruginosa* attachment to epithelial cells and delays infection-induced cytotoxicity

Because surface attachment contributes to both persistence on abiotic materials and host colonization, we next asked whether dalberoside could limit bacterial retention on mammalian cell surfaces and alters infection-associated outcomes. We first incubated mNeonGreen-expressing *P. aeruginosa* with dalberoside or vehicle on Madin–Darby canine kidney (MDCK) epithelial monolayers, then washed the monolayers and quantified retained bacteria by confocal (Fig. 4a).^27^ Dalberoside significantly reduced cell-associated bacterial signal (Fig. 4b, c), demonstrating that its anti-attachment activity extends to mammalian cell surfaces. To determine whether this effect influenced infection-associated host-cell survival, we next turned to a model amenable to sensitive, continuous monitoring of cytotoxicity, for which MDCK was not suitable.

**Fig. 4.**
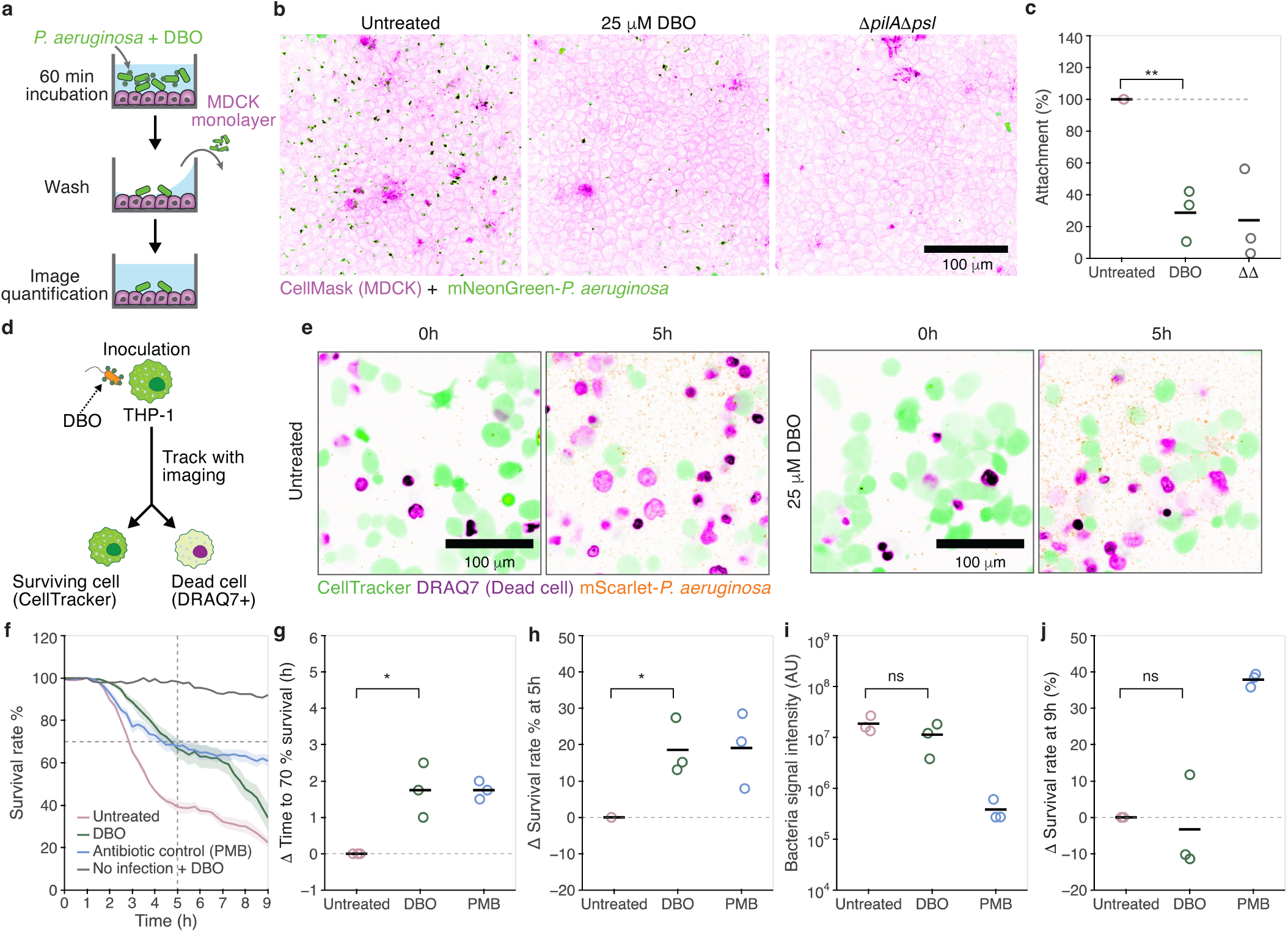
Dalberoside (DBO) reduces *Pseudomonas aeruginosa* attachment to epithelial cells and delays infection-induced THP-1 cytotoxicity. **a,** MDCK attachment assay workflow. mNeonGreen-expressing *P. aeruginosa* was incubated with CellMask-stained MDCK monolayers for 60 min with DMSO or 25 µM DBO before washing and imaging. **b,** Representative images of bacterial attachment following treatment with DMSO or DBO; Δ*psl*Δ*pilA* served as an attachment-defective control. MDCK membranes are shown in magenta and bacteria in green. **c,** Bacterial attachment normalized to the DMSO-treated wild-type control. **d,** THP-1 infection assay workflow. Differentiated THP-1 cells were infected with mScarlet-expressing *P. aeruginosa* with DMSO or 25 µM DBO; polymyxin B (PMB; 2 µg/ml) was added at 1 h post-infection (hpi). Uninfected DBO-treated cells served as a cytotoxicity control. **e,** Representative images at 0 and 5 hpi. CellTracker-labeled THP-1 cells are shown in green, DRAQ7-positive cells in magenta and bacteria in orange. **f,** Representative THP-1 survival time courses; lines and shading indicate the mean ± SEM of technical replicate wells. **g,** Change in time to 70% survival relative to matched DMSO controls. **h,j,** Change in survival at 5 hpi (**h**) and 9 hpi (**j**). **i,** Integrated bacterial mScarlet signal at 5 hpi. Positive Δ values in **g,h,j** indicate delayed cytotoxicity or increased survival relative to DMSO. Colored open circles represent independent biological replicates and black horizontal lines indicate the mean; *n* = 3. Statistical analyses are described in Methods. *P* < 0.05, *; *P* < 0.01, **; *P* < 0.001, ***; *P* < 0.0001, ****; ns, not significant. Scale bars, 100 µm.

To determine whether dalberoside conferred protection to host cells against *P. aeruginosa* virulence, we used THP-1-derived macrophages as an infection model.^28^ We infected differentiated THP-1 cells with mScarlet-expressing *P. aeruginosa* premixed with 25 µM dalberoside or vehicle, and monitored host-cell survival over 9 h by confocal microscopy (Fig. 4d, e). The compound did not affect the viability of uninfected THP-1 cells at this concentration (Fig. 4f and Extended Data Fig. 5). Dalberoside delayed *P. aeruginosa*-induced THP-1 cell death relative to the vehicle control (Fig. 4f, g). At 5 h, dalberoside increased THP-1 survival by approximately 20% relative to the vehicle control, similar to the observation with the antibiotic positive control, polymyxin B (PMB) (Fig. 4h). Whereas PMB reduced bacterial fluorescence, consistent with its bactericidal activity, dalberoside did not measurably reduce bacterial abundance (Fig. 4i). By the experimental endpoint, dalberoside no longer conferred detectable protection. This transient effect is consistent with an anti-virulence mechanism that delays, rather than prevents, cytotoxicity without limiting bacterial accumulation in the confined assay environment (Fig. 4j).

### Dalberoside targets the bacterial outer membrane

The rapid onset and broad efficacy of dalberoside across attachment contexts, together with its lack of bactericidal activity, suggest that it acts on a shared determinant of bacterial surface association. To test whether dalberoside instead acted through specific adhesion machinery, we examined its activity in mutants deficient in two major *P. aeruginosa* attachment factors. Dalberoside was already against Δ*psl* in screening conditions. It also inhibited attachment of a Δ*pilA* mutant lacking the major subunit of type IV pili (T4P) involved in early surface colonization (Extended Data Fig. 6). Thus, no known specific attachment structure is required for activity. Together with its amphipathic structure and rapid action, these results suggest that dalberoside might instead alter physicochemical properties of the bacterial envelope.

We therefore asked whether dalberoside associates with the bacterial envelope. We generated a fluorescent analogue by conjugating BODIPY to the glycan headgroup while leaving the lipid tail unmodified (Fig. 5a).^25^ In both *P. aeruginosa* and *A. baumannii*, fluorescence from the labelled analogue was enriched at the cell periphery and spatially overlapped with the signal from the membrane dye FM4-64 (Fig. 5b, c, Extended Data Fig. 7). These observations support association of the labeled dalberoside analogue with the bacterial envelope.

**Fig. 5.**
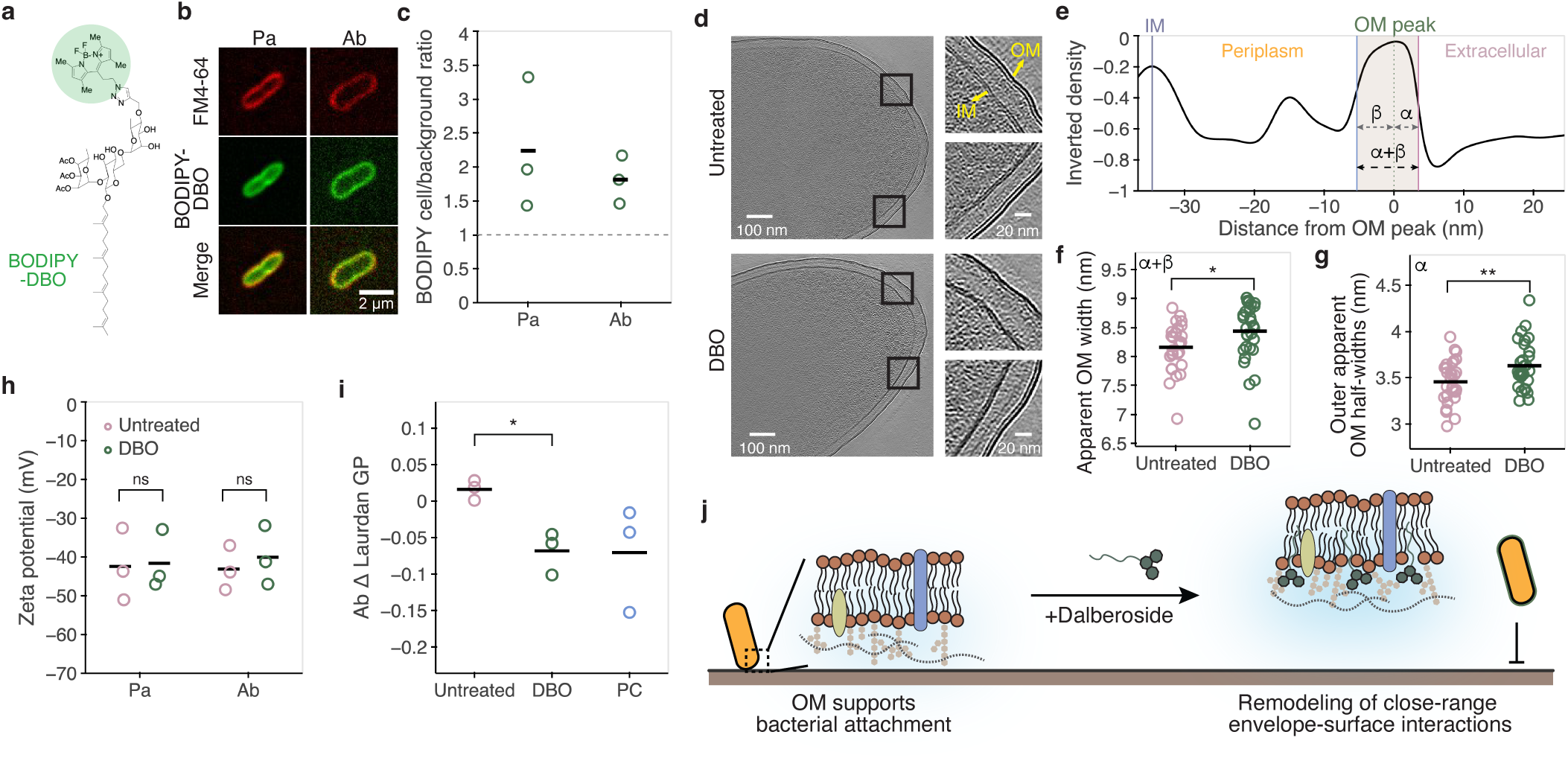
Dalberoside (DBO) associates with bacterial envelopes and alters outer membrane interfacial properties. **a,** Structure of the BODIPY-labeled DBO analog. **b,** Representative images of *P. aeruginosa* at 45 min and *A. baumannii* at 30 min after exposure to BODIPY–DBO under an agarose pad. FM4-64 and BODIPY–DBO are shown in red and green, respectively. Scale bars, 2 µm. **c,** Cell-associated BODIPY–DBO fluorescence at 60 min relative to the mean field background outside cell-associated regions. The dashed line at 1 indicates no enrichment over background. **d,** Representative computational slices of cryo-electron tomograms of *P. aeruginosa* treated with DMSO or 25 µM DBO. **e,** Outer membrane density profile analysis. Profiles were aligned to the outer membrane density peak, and apparent width was defined by the outer (extracellular side) and inner (periplasm side) positions of maximal absolute density change. α and β denote the distances from the peak to the outer and inner edges, respectively. **f,g,** Apparent outer membrane width (**f**) and outer-facing half-width α (**g**). DMSO, *n* = 31 cells; DBO, *n* = 30 cells. Cells were sampled from one independently prepared specimen per condition. **h,** Zeta potential of *P. aeruginosa* treated with DMSO or 25 µM DBO and *A. baumannii* treated with DMSO or 12.5 µM DBO. **i,** Change in Laurdan generalized polarization (ΔGP) in *A. baumannii* 10 min after treatment with DMSO, 12.5 µM DBO or 0.5% benzyl alcohol as a positive control, relative to the pretreatment baseline. **j,** Proposed model for DBO-mediated inhibition of bacterial attachment through remodeling of the close-range envelope-surface interaction. In **c,h,i**, colored open circles represent independent biological replicates and black horizontal lines indicate the mean; *n* = 3. Panel **b** shows representative images from one of three independent biological replicates. Statistical analyses are described in Methods. *P* < 0.05, *; *P* < 0.01, **; *P* < 0.001, ***; *P* < 0.0001, ****; ns, not significant.

To resolve the effects of unmodified dalberoside on the bacterial envelope at higher spatial resolution, we used *in situ* cryo-electron tomography (cryo-ET). Both vehicle- and dalberoside-treated *P. aeruginosa* cells retained intact outer membranes and inner membranes, with no evidence of gross envelope disruption (Fig. 5d). Quantification of outer-membrane-centered density profiles nevertheless revealed a significant increase in apparent outer membrane width following dalberoside treatment (Fig. 5e, f and Extended Data Fig. 8a, b). The outer-facing part of the outer membrane mainly contributed to this increase, whereas the inner-facing part remained unchanged (Fig. 5g and Extended Data Fig. 8c). This asymmetric broadening is consistent with dalberoside-induced structural remodeling of the outer-leaflet of the *P. aeruginosa* outer membrane.

Having identified dalberoside-induced structural remodeling of the *P. aeruginosa* outer membrane, we next asked whether it was accompanied by changes in cell-surface physicochemical properties relevant to attachment. We could not detect a change in zeta-potential in either *P. aeruginosa* or *A. baumannii*, indicating that dalberoside does not substantially alter surface electrostatics (Fig. 5h).^20^ We then used Laurdan staining to probe the local membrane interfacial environment. Because of its pronounced detachment phenotype and compatibility with Laurdan measurements, we performed this analysis in *A. baumannii*. Dalberoside treatment reduced Laurdan generalized polarization (GP) relative to the vehicle control, consistent with increased membrane-interfacial polarity and hydration and potentially reduced lipid packing (Fig. 5i).^29^ Together, the asymmetric outer membrane broadening observed in *P. aeruginosa* and the altered membrane interfacial properties detected in *A. baumannii* support a model in which dalberoside remodels the bacterial envelope without gross membrane disruption. These envelope-associated changes may gently alter the short-range interactions required for stable bacterial attachment.

## Discussion

We identified dalberoside as a non-bactericidal inhibitor of bacterial surface attachment. Dalberoside is a natural glycolipid isolated from *Dalbergia oliveri* whose biological activity has not been previously characterized.^30,31^ Dalberoside inhibited attachment and biofilm-associated phenotypes in the Gram-negative pathogens *P. aeruginosa* and *A. baumannii* without detectable effects on bacterial viability or membrane permeabilization. These findings demonstrate that bacterial surface association can be chemically perturbed independently of growth inhibition.

Our findings support a working model in which dalberoside non-disruptively remodels the Gram-negative cell-envelope interface (Fig. 5j). We propose that dalberoside inserts into the outer-leaflet of the outer membrane via it hydrophobic tail, orienting its glycan headgroup remains towards the aqueous extracellular environment. Such an arrangement could alter local lipid organization, hydration or other short-range interactions required for stable bacterium–surface contact.^20^ Related observations support this mechanism. Geranylgeranoic acid, which resembles the lipid moiety of dalberoside, reduces *Staphylococcus aureus* adhesion at sub-microbicidal concentrations.^32^ Hydrated polysaccharide-rich surface structures can restrict close cellular contact through steric and interfacial effects,^33^ and rhamnolipid biosurfactants produced by *P. aeruginosa* can weaken bacterial adhesion by altering cell-surface properties.^34^ Rather than causing classical surfactant-like membrane disruption, dalberoside altered outer membrane architecture and interfacial properties without measurably affecting membrane integrity, surface charge, growth, or viability. These findings suggest that subtle remodeling of the outer membrane interface is sufficient to disrupt bacterial attachment.

The consequences of this remodeling may depend on species-specific envelope organization and surface-association strategies. *P. aeruginosa* uses cellular appendages to actively explore surfaces and possesses O-antigen-containing lipopolysaccharide, which forms an extended layer at the cell surface.^35,36^ By contrast, the *A. baumannii* outer membrane contains lipooligosaccharide lacking an extended O-antigen.^37^ These architectural differences could influence dalberoside access, partitioning or the sensitivity of envelope–surface interactions to interfacial remodeling. They may therefore contribute to the species-dependent differences between inhibition of initial attachment and detachment of surface-associated bacteria.

The effects of dalberoside extended beyond attachment to abiotic surfaces. Dalberoside reduced *P. aeruginosa* retention on epithelial cells and transiently delayed infection-associated cytotoxicity without measurably reducing bacterial abundance. In *A. baumannii*, dalberoside’s ability to detach surface-associated cells and reduce preformed biofilms motivates investigations on clinically relevant materials susceptible to bacterial colonization and biofouling.^26^ Together, our results provide initial evidence that perturbing attachment can influence both biofilm-associated persistence and early host–pathogen interactions in the absence of conventional antibacterial activity. They also support further evaluation of dalberoside in settings where surface association and biofilm formation contribute to disease, such as wound infections,^38^ and as a potential adjuvant to antibiotics against surface-associated populations.^39,40^

Overall, we identified the Gram-negative outer membrane interface as a chemically addressable determinant of bacterial surface attachment. Dalberoside provides an alternative to adhesin-specific strategies, which may be difficult to generalize across bacterial species and surface environments. More broadly, our study demonstrates the value of image-based phenotypic screening for discovering molecules that alter bacterial colonization rather than growth and may therefore be overlooked by conventional antibacterial screens.

## Materials and Methods

### Bacterial strains and culture conditions

*Pseudomonas aeruginosa* PAO1 ATCC 15692 (American Type Culture Collection) and the derivative mutant strains were used. The construction of the *P. aeruginosa* in-frame deletion mutants (Δ*pilA*, Δ*pslBCD*, Δ*pilA*Δ*pslBCD*, Δ*popB*) and fluorescent strains constitutively expressing mScarlet or mNeonGreen was previously described.^28,41,42^ *Acinetobacter baumannii* ATCC 19606 was included as the WT strain. Unless otherwise specified, bacteria were cultured in lysogeny broth (LB, Roth) at 37 °C with shaking at 220 rpm. LB agar plates were prepared by adding 1.5 % agar (Fisher Scientific) to LB medium and gentamicin (Sigma-Aldrich) was added as required for fluorescent strains. Overnight cultures were diluted 1:100 and grown to the optical densities specified for each assay.

### Compounds

Dalberoside was purchased from AnalytiCon Discovery, prepared as a 10 mM stock in DMSO (Fisher Bioreagents) and stored at −20 °C. Working solutions were diluted in the assay medium, with matched final DMSO concentrations in all conditions. Chemically synthesized dalberoside and the BODIPY-conjugated dalberoside analog 14-CWA-019 were provided by the collaborating laboratory; their synthesis and characterization are described in reference.^25^

### Image-based attachment and detachment assays

A library of 2,627 purified natural products was screened at 30 µM in PhenoPlate 96-well microplates (Revvity). The library comprised compounds derived from fractionated plant and bacterial extracts from commercial libraries of Analyticon Discovery GmbH (Germany) and InterBioScreen Ltd. (Russia) and prepared as 10 mM DMSO stocks by the Biomolecular Screening Facility at EPFL, Lausanne. *P. aeruginosa* cultures were grown to OD₆₀₀ = 1.5, pelleted, and resuspended to OD₆₀₀ = 1.0 with LB. Bacteria were added to microplates at 50 µl and incubated with compounds for 30 min. Wild-type (WT) cells were washed to remove non-attached cells before imaging, whereas Δ*psl* cells were imaged without washing. Assay performance was benchmarked using WT and Δ*psl* cells under post-wash conditions and Δ*psl* and Δ*psl*Δ*pilA* cells under pre-wash conditions; both assay configurations yielded Z′ factors >0.8. Attachment was normalized to the corresponding DMSO control, and compounds that reduced attachment by >50% in both configurations were selected as primary hits.

For subsequent concentration–response assays, *P. aeruginosa* and *A. baumannii* cultures were grown to OD₆₀₀ = 1.5 and 1.0, respectively. Bacteria were pelleted and resuspended to OD₆₀₀ = 1.0 with LB. Bacteria cultures (50 µl per well) were incubated with the indicated concentrations of dalberoside for 30 min and imaged after washing, except for *P. aeruginosa* Δ*psl*, which was imaged without washing. For detachment assays, bacteria were allowed to attach for 30 min and imaged after removal of non-attached cells. Attached bacteria were then treated with dalberoside or DMSO for 30 min, washed and imaged again. Detachment was calculated from the reduction in segmented cell area relative to the initial attachment measured in the same well.

Phase-contrast images were acquired using a Nikon ECLIPSE Ti epifluorescence microscope equipped with a Hamamatsu ORCA-flash4.0 camera, a ×40 Plan Apo phase-contrast objective (NA 0.95) and NIS-Elements software (version AR 5.02). Uneven illumination was corrected computationally, and bacteria were segmented by intensity thresholding. Surface attachment was quantified as the total segmented cell area per image.

### Growth and CFU assays

Bacteria were grown to OD₆₀₀ = 0.5 – 0.8 in exponential phase and were adjusted to OD₆₀₀ = 0.3. The cultures were then diluted 200–fold in Mueller–Hinton broth (MHB) or LB. Bacteria were dispensed with dalberoside or matched DMSO into 384-well plates (Corning, 3701; 50 µl per well) and incubated at 37 °C in a Spark plate reader (Tecan). OD₆₀₀ was monitored for 20 h. The minimum inhibitory concentration was defined as the lowest concentration preventing detectable growth.^43^ The area under each growth curve (AUC) from 0–20 h was calculated for each well by trapezoidal integration. AUC values were normalized to the mean vehicle-control AUC within each biological replicate, after which technical replicate wells were averaged. For viable-cell-count measurements, cultures prepared in LB were treated with dalberoside or DMSO for 24 h at 37 °C with shaking at 220 rpm. Samples were serially diluted in PBS (BioConcept), and 5 µl aliquots were plated on LB agar plates and incubated overnight before CFU/ml was determined.

### Propidium iodide membrane-integrity assays

Membrane integrity was determined using the propidium iodide (PI) assay.^44^ For bulk measurements, exponential-phase cells were resuspended in LB to OD₆₀₀ = 0.7 and treated with DMSO, dalberoside or polymyxin B (Apollo Scientific) in the presence of 10 µg/ml propidium iodide (PI; Cayman Chemical, 10008351). Samples were analyzed in technical duplicate in PhenoPlate 96-well plates using a Spark reader at 37 °C for 2 h. PI fluorescence was measured at 493 nm excitation and 636 nm emission. Fluorescence was divided by OD₆₀₀ and normalized to the initial value in each well.

For single-cell analysis, cells were treated for 30 min with DMSO, dalberoside or polymyxin B in the presence of 6.25 µg/ml PI. Samples were immobilized under 1% agarose–PBS pads and imaged using a Nikon ECLIPSE Ti microscope/Hamamatsu ORCA-Flash4.0 camera, a ×100 Plan Apo λ phase-contrast oil objective (NA 1.45) and TxRed-A-Basic-NTE filters. Cells were segmented from phase-contrast images, and background-corrected PI fluorescence was measured per cell. Cells with mean PI fluorescence >2,000 arbitrary units were classified as PI-positive.

### Biofilm assays

The biofilm assay was adapted from previous publications.^45,46^ For biofilm formation, exponential-phase *P. aeruginosa* and *A. baumannii* cultures were adjusted to OD₆₀₀ = 0.02 and 0.1, respectively. Cells were incubated with dalberoside or DMSO in PhenoPlate 96-well plates (100 µl/well) at 37 °C for 24 h under static, humidified conditions. After non-associated cells being removed, the wells were washed with PBS and dried at 60 °C for 30 min. Biofilms were stained with 200 µl of 0.1% crystal violet for 10 min, washed and air-dried. Bound dye was solubilized in 200 µl of 95% ethanol and measured at 595 nm using a Spark reader with SparkControl software. Biofilm biomass was normalized to the matched DMSO control.

For CFU-based distribution analysis, supernatant and well-associated populations were recovered separately after 24 h. Well-associated populations were recovered by repeated pipetting (>10 times). Both fractions were serially diluted and plated. Total recoverable CFUs were calculated as their sum, and the well-associated population was expressed as a percentage of the total.

For preformed-biofilm reduction, biofilms were established for 23 h from cultures adjusted to OD₆₀₀ = 0.1. Planktonic cells were removed and preformed biofilms were treated with 50 µl of fresh LB containing dalberoside or DMSO for 1 h before crystal violet quantification.

### Microfluidic flow-based detachment

Three-channel PDMS (SYLGARD 184, Dow Corning, base-to-curing-agent ratio 10:1) chips were fabricated as previously described.^47^ Each channel was 1 cm long, 500 µm wide and 50 µm deep, with 1-mm-diameter inlet and outlet ports and no bubble trap. The PDMS devices were reversibly sealed to glass coverslips without plasma treatment, forming a sufficiently robust seal for perfusion, and connected to BTPE-20 polyethylene tubing (Instech) with a syringe pump (KD Scientific).

Exponential-phase bacterial cultures were adjusted to OD₆₀₀ = 0.6 in LB and introduced into the channels. *P. aeruginosa* and *A. baumannii* were allowed to attach statically for 30 min before perfusion with LB containing DMSO or dalberoside. Flow rates were 0.5 µl/min for *P. aeruginosa* and 1 µl/min for *A. baumannii*.

Phase-contrast images were acquired at 1-min intervals using the Nikon ECLIPSE Ti2/Prime 95B system and ×40 Plan Apo phase-contrast objective (NA 0.95). *P. aeruginosa* and *A. baumannii* were monitored for 60 and 30 min, respectively. Segmented cell area was normalized to the first frame, and endpoint retention was quantified at the final time point.

### MDCK attachment assay

MDCK attachment assay was adapted from previous studies.^27,48^ MDCK cells (RRID: CVCL_0422) were maintained in DMEM (Gibco) supplemented with 5% fetal bovine serum (FBS, Thermo Fisher) at 37 °C with 5% CO_2_. Cells were seeded at 1 × 10⁴ cells per well in PhenoPlate 96-well plates and cultured for 3 d to form highly confluent monolayers. Two hours before infection, membranes were labeled with CellMask Deep Red dye (Life Technologies, 2.5 µg/mL). During infection, medium was replaced with phenol red-free DMEM (Gibco) containing 3% fetal bovine serum. The mNeonGreen-expressing *P. aeruginosa* was added at an estimated MOI of 5 with DMSO or dalberoside. The mNeonGreen-expressing Δ*psl*Δ*pilA* strain served as an attachment-defective control. After 60 min at 37 °C, monolayers were washed three times with PBS.

Images were acquired using a Nikon Eclipse Ti2 microscope/Yokogawa CSU-W1 spinning-disk /Prime 95B camera and a ×40 water-immersion objective (NA 1.15). Brightfield, GFP and Cy5 images were collected as 25-plane z-stacks at 0.5-µm intervals. GFP stacks were converted to maximum-intensity projections, background-corrected and segmented. Attachment was quantified as GFP-positive area per field and normalized to the DMSO-treated wild-type control within each biological replicate.

### THP-1 infection and cytotoxicity assay

THP-1 cells (ATCC TIB-202) were maintained in RPMI 1640 containing 10% FBS and 1× GlutaMAX (all from Thermo Fisher). Cells were seeded at 20,000 cells per well and differentiated with 100 nM phorbol 12-myristate 13-acetate (PMA, Adipogen) for 3 days.

The THP-1 infection protocol was performed as previously described.^28^ Cells were labeled with 1 µM CellTracker Green CMFDA (Thermo Fisher) for 30 min. Medium was then replaced with phenol red-free RPMI containing 10% FBS, 1 µM cytochalasin D (Cayman Chemical) and 3 µM DRAQ7 (Thermo Fisher).^49^ After 10 min, the mScarlet-expressing *P. aeruginosa* was added at an MOI of 10 with DMSO or dalberoside. Polymyxin B, used as antibiotic control, was added at 1 h post-infection. Uninfected cells treated with dalberoside were included as a cytotoxicity control.

Single-plane images were acquired every 15 min for 9 h using the Nikon Eclipse Ti2 equipped with Yokogawa CSU-W1 spinning-disk, Prime95B camera, and a ×20 Apo objective (NA 0.75). Initial CellTracker-positive counts defined the total cell population, and DRAQ7-positive objects were detected at each time point. Survival was calculated relative to the initial cell number. Bacterial abundance was quantified from segmented mScarlet area and integrated fluorescence. Technical wells were averaged within each biological replicate. Survival endpoints, time to 70% survival and bacterial signal were extracted for statistical analysis.

### BODIPY-labeled dalberoside association

Exponential-phase bacteria were stained with 5 µM FM4-64FX (Invitrogen) for 15 min. Cells were washed, resuspended in PBS to OD₆₀₀ = 0.6 and immobilized under PBS pads containing 0.8% agarose (Roth). A 4-µl volume of 25 µM BODIPY-labeled dalberoside was applied to the opposite side of the pad. Images were acquired every minute for 60 min using the Nikon Eclipse Ti2 equipped with Yokogawa CSU-W1 spinning-disk, HAMAMATSU ORCA-flash4.0 camera and a ×100 oil-immersion objective (NA 1.45). BODIPY and FM4-64 signals were collected using the GFP and Cy3 channels. BODIPY fluorescence within bacterial masks was divided by the mean field background measured outside buffered cell-associated regions. Association was quantified over time and at 60 min.

### Cryo-electron tomography

*P. aeruginosa* Δ*popB* mutant, a low virulence strain, was used in this assay. Overnight culture was subcultured in LB media and grown to OD_600_ = 0.3. Before vitrification, cells were incubated for 30 minutes with either 25 μM dalberoside or a matched DMSO vehicle control. BSA gold tracers (6 nm, Electron Microscopy Sciences) were added to the samples prior vitrification in a 1:5 volume ratio. 4 μl of bacterial cultures were applied onto Quantifoil R2/2 200 mesh copper grids, previously glow discharged on both sides for 25 s at 15 mA using PELCO easiGlow™ machine. Vitrification was done on a Leica EM GP2 automatic plunger (Thermo Fisher Scientific) at 95% humidity with a waiting time of 10 s and back-side blotting of 3 s. Grids were plunge-frozen into liquid ethane and stored under liquid nitrogen until imaging.

All imaging was done on a transmission electron microscope E-CFEG Titan Krios G4 (Thermo Fisher Scientific) operated at 300 kV, equipped with a Selectris X energy filter and a Falcon 4i direct detector at the Dubochet Center for Imaging (**DCI,** University of Lausanne). Data collection was acquired using Tomo5 software (Thermo Fisher Scientific) at the nominal magnification of 42,000**x,** giving a calibrated pixel size of 3.06 Å. Tilt series were collected using a range of -3.0 to -7.0 μm nominal defoci with a 0.5 μm steps and a total dose of 130 e-**/**Å^2^ following a dose-symmetric tilt scheme^50^ from -60° to 60° in 3° steps. All collected tilt**-**series were first pre**-**processed using on**-**the**-**fly Scipion tomogram reconstruction pipeline^51^ available at the **DCI.** Tilt**-**series were first motion corrected using MotionCor2^52^ and subsequently processed using the IMOD software package^53,54^ integrated into SCIPION framework, and cryoNAV^55^ was used to refine fiducial tilt-series alignment. CTF estimation was done using AreTomo^56^ inside SCIPION. Weighted back projected reconstructed tomogram were binned 4x and denoised in cryo-CARE^57^ prior analysis. For membrane measurements among conditions, ten central computational slices were summed projected employing customized MATLAB scripts in the software package *Dynamo*^58^. Summed projections were analyzed using a custom Python pipeline. Candidate outer membrane contours were identified by automated image enhancement, ridge detection and skeletonization, followed by manual review to exclude incorrectly segmented regions. One-dimensional density profiles normal to the retained outer membrane contours were extracted, aligned to the outer membrane density peak and averaged for each cell. Outer membrane edge positions were defined by local maxima in the absolute gradient of the averaged density profile.

### Zeta-potential measurement assay

Zeta-potential was measured using 900-µl samples in DTS1070 cuvette (Malvern) to quantify the bacterial surface charge.^59^ Exponential-phase bacteria were washed twice with Milli-Q water and resuspended in Milli-Q water to OD₆₀₀ = 0.3. *P. aeruginosa* and *A. baumannii* were treated with dalberoside or DMSO for 30 min directly in the cuvette. Zeta potential was measured at 25 °C using a Zetasizer Nano ZS (Malvern).

### Laurdan generalized polarization assay

Exponential-phase *A. baumannii* was labeled with 10 µM Laurdan for 20 min at 37 °C in the dark. Cells were washed three times with pre-warmed PBS and resuspended to OD₆₀₀ = 0.8. Samples were transferred to PhenoPlate 96-well plates at 100 µl per well. Fluorescence was measured at 37 °C using 350 nm excitation and emission at 460 and 500 nm. After a 5-min baseline, DMSO, dalberoside or 0.5% benzyl alcohol (Sigma-Aldrich) was added and measurements continued for 30 min at 1-min intervals. Generalized polarization was calculated as

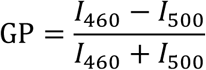

ΔGP was defined as the GP value 10 min after treatment minus the pretreatment baseline.

### Statistical analysis

Statistical analyses were performed in Python. Unless otherwise specified, technical replicates were averaged within each biological replicate, and biological-replicate means were used for statistical inference. CFU and THP-1 bacterial mScarlet measurements were log10-transformed before analysis, whereas percentage outcomes were analyzed on their original scale. Tests were two-sided unless a directional hypothesis was specified a priori. Positive controls were included for assay validation and analyzed descriptively. Statistical significance was defined as (P < 0.05); Holm-adjusted P values were used for families of concentration-specific comparisons.

Screen performance was evaluated using the Z′ factor, and compound-screening results were analyzed descriptively. Attachment and static-biofilm concentration–response data were fitted to biological-replicate means using four-parameter log-logistic models. Growth, bulk PI, Laurdan, THP-1 survival and BODIPY–dalberoside fluorescence time courses were analyzed descriptively.

For growth AUC, values normalized to the within-replicate vehicle mean were compared with 100% using two-sided one-sample t-tests, with Holm adjustment across concentrations within each bacterial strain and medium. Uninfected THP-1 viability was compared with matched vehicle controls using two-sided paired t-tests with Holm adjustment across concentrations. Static-biofilm formation at 25 µM was compared with 100% using a two-sided one-sample t-test. Preformed-biofilm biomass was compared with 100% using one-sided one-sample t-tests, based on the prespecified hypothesis that dalberoside reduces preformed-biofilm biomass.

Matched dalberoside and vehicle conditions for endpoint CFU, image-based PI-positive fraction, static-biofilm CFU distribution, plate-based and microfluidic detachment, THP-1 bacterial mScarlet signal, zeta potential and Laurdan ΔGP were compared using two-sided paired t-tests. MDCK attachment was compared with 100% using a one-sided one-sample t-test based on the prespecified hypothesis that dalberoside reduces bacterial attachment. THP-1 survival endpoints and time to 70% survival were analyzed using one-sided paired t-tests based on the prespecified hypothesis that dalberoside delays infection-induced host-cell death. BODIPY–dalberoside envelope-association measurements were summarized descriptively.

Cryo-electron tomography measurements were obtained from one specimen per condition. Individual cells were therefore treated as observational units in exploratory two-sided Welch’s t-tests and were not interpreted as biological replicates.

## Acknowledgments

We thank Marc Chambon, Julien Bortoli Chapalay and Antoine Gibelin from the Biomolecular Screening Facility at EPFL for assistance with assay development and compound-library preparation. We thank Tania Distler for developing the THP-1 cell assay protocol used in this study, and Sourabh Monappa and Alice Cont for designing the microfluidic chip used in this study. We thank the Molecular and Hybrid Materials Characterization Center at EPFL for assistance with Zetasizer Nano ZS measurements. We thank all members of the Dubochet Center for Imaging for maintaining and providing access to high-end cryo-EM equipment. This work was supported by the Swiss National Science Foundation through the National Centre of Competence in Research AntiResist (grant nos. 51NF40-180541 and 51NF40-225154), the Swiss National Science Foundation (grant no. 310030-189084) and the Fondation Jacqueline et Virginie Beytout to A.P.; SNSF Starting Grant (TMSGI3_218251) and the Foundation Pierre Mercier pour la Science to P.P.N.; Ministry of Education Singapore, MOE-T2EP30223-0002, MOE-T2EP10124-0001, and Tier 1 grant RG2/25, and the National Medical Research Council, MOH-002082, to Y.Q. ChatGPT-5 was used to develop analysis code, polish grammar and syntax in this manuscript.

## Author information

### Contributions

Conceptualization: H.Y.H. and A.P.

Data curation: H.Y.H., P.P.N. and E.N.A.-S.

Formal analysis: H.Y.H.

Funding acquisition: A.P., P.P.N. and Y.Q.

Investigation: H.Y.H.

Methodology: H.Y.H., P.P.N., E.N.A.-S. and C.A.

Project administration: H.Y.H.

Supervision: A.P.

Visualization: H.Y.H.

Writing-original draft: H.Y.H. and A.P.

Writing-review and editing: H.Y.H., A.P., P.P.N and E.N.A.-S.

Corresponding authors: A.P.

## Extended Data

**Extended Data Fig. 1.**
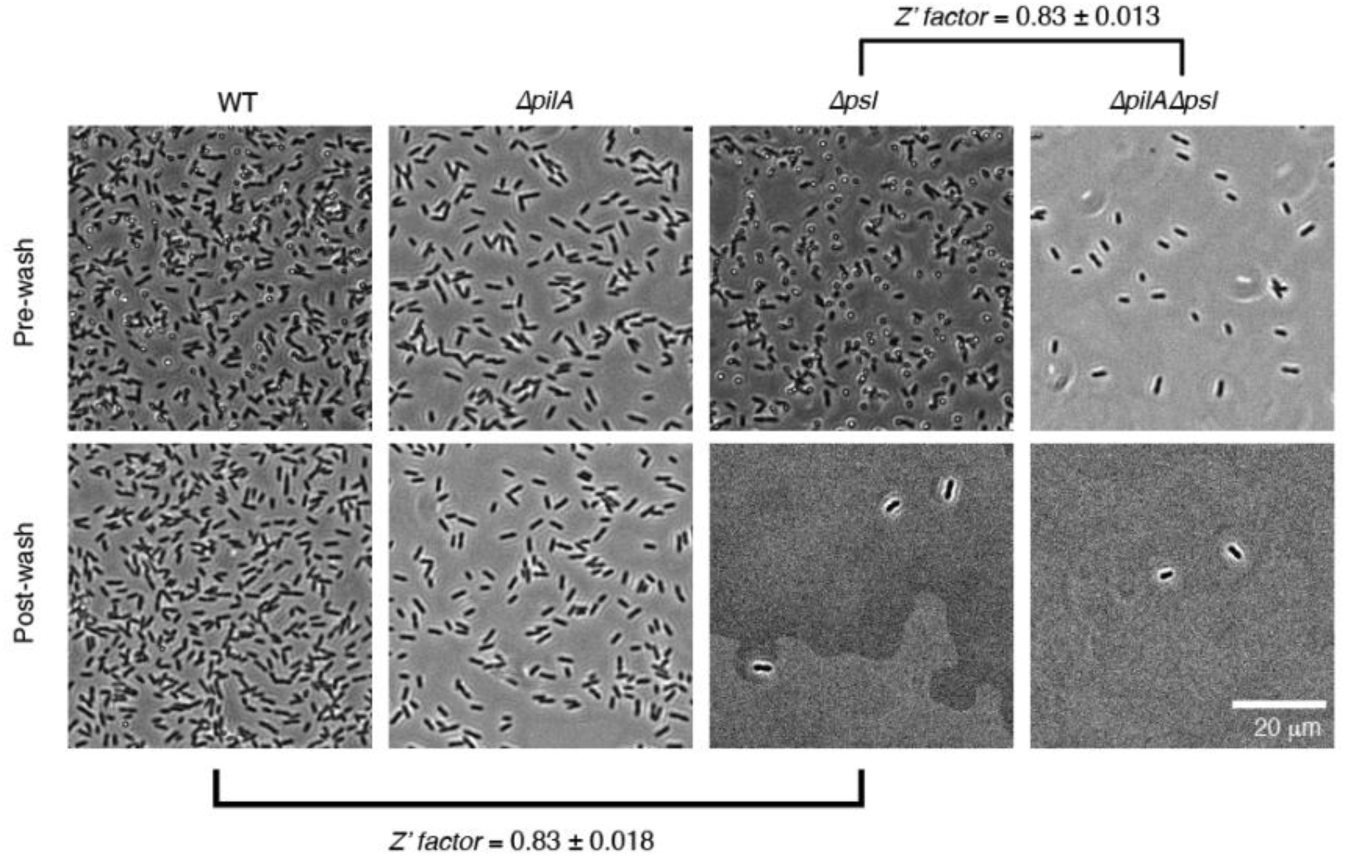
Benchmarking attachment phenotypes in *P. aeruginosa* attachment-defective mutants. Representative images of WT, Δ*psl*, Δ*pilA* and Δ*psl*Δ*pilA* cells after 30 min of surface attachment, imaged before or after washing to remove non-attached cells. Z′ factors were calculated between WT and Δ*psl* under post-wash conditions and between Δ*psl* and Δ*psl*Δ*pilA* under pre-wash conditions to select the screening configurations. Scale bars, 20 µm.

**Extended Data Fig. 2.**
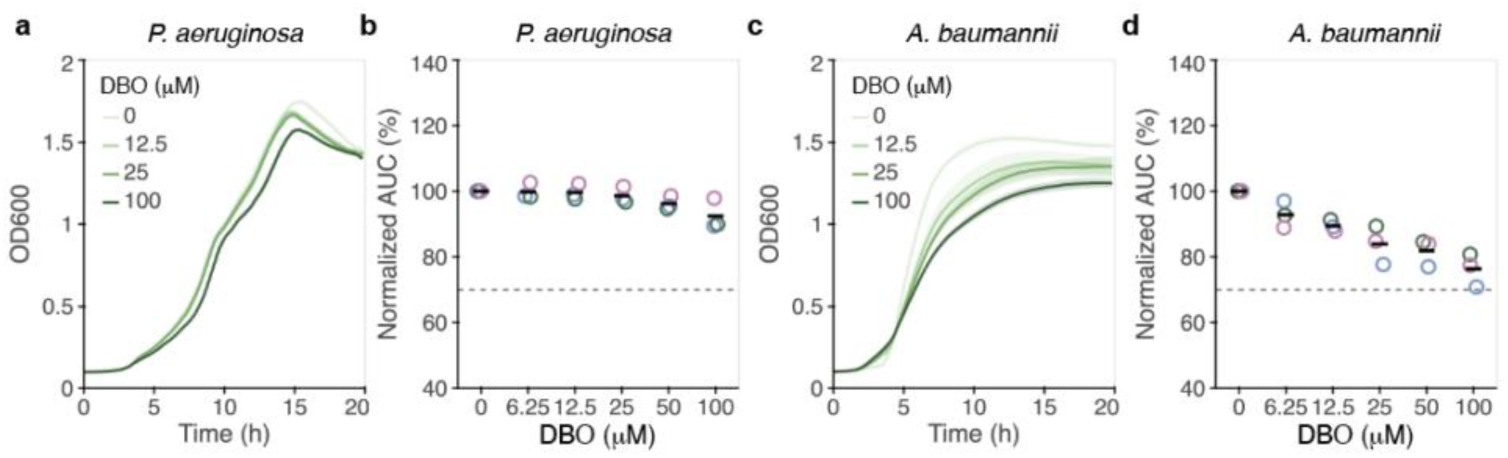
Growth of *P. aeruginosa* and *A. baumannii* during dalberoside (DBO) treatment in LB. **a–d,** Representative growth curves (**a,c**) and normalized area-under-the-curve (AUC) analyses (**b,d**) of *P. aeruginosa* (**a,b**) and *A. baumannii* (**c,d**) treated with the indicated DBO concentrations for 20 h. Growth was monitored by OD₆₀₀. AUC values were normalized to the mean DMSO control within each biological replicate. In **b,d**, colored open circles represent independent biological replicates and black horizontal lines indicate the mean; *n* = 3. Panels **a,c** show one representative experiment from three biological replicates. Statistical analyses are described in Methods. *P* < 0.05, *; *P* < 0.01, **; *P* < 0.001, ***; *P* < 0.0001, ****; ns, not significant.

**Extended Data Fig. 3.**
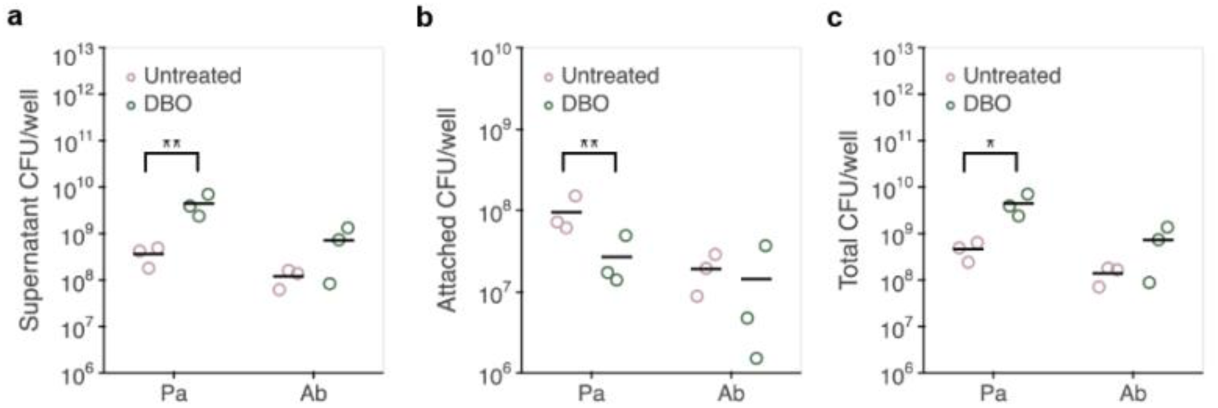
CFU-based distribution of *P. aeruginosa* and *A. baumannii* after 24 h static biofilm formation with DMSO or 25 µM dalberoside (DBO). **a,** Supernatant CFUs recovered per well. **b,** Well-associated CFUs recovered after removal of the supernatant. **c,** Total recoverable CFUs per well, calculated as the sum of the supernatant and well-associated fractions. Colored open circles represent independent biological replicates and black horizontal lines indicate the mean; *n* = 3. Statistical analyses are described in Methods.

**Extended Data Fig. 4.**
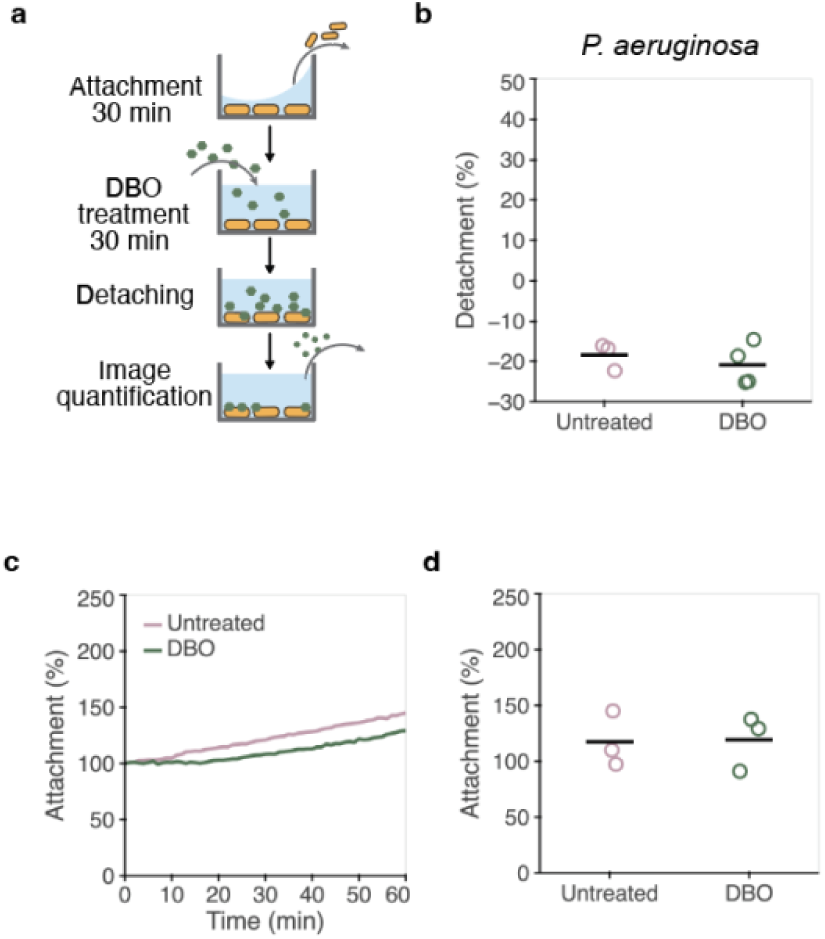
Dalberoside (DBO) does not promote detachment of surface-associated *P. aeruginosa*. **a,** Plate-based detachment assay workflow. Bacteria were allowed to attach for 30 min and treated with DMSO or 25 µM DBO for 30 min before washing and reimaging. **b,** Detachment calculated from paired pre- and post-treatment segmented cell areas within each well. **c,d,** Representative time course (**c**) and endpoint quantification (**d**) of surface-associated *P. aeruginosa* during microfluidic perfusion with DMSO or 25 µM DBO. Values were normalized to the initial time point; the endpoint was measured after 60 min. In **d**, colored open circles represent independent biological replicates and the black horizontal line indicates the mean; n = 3. Statistical analyses are described in Methods.

**Extended Data Fig. 5.**
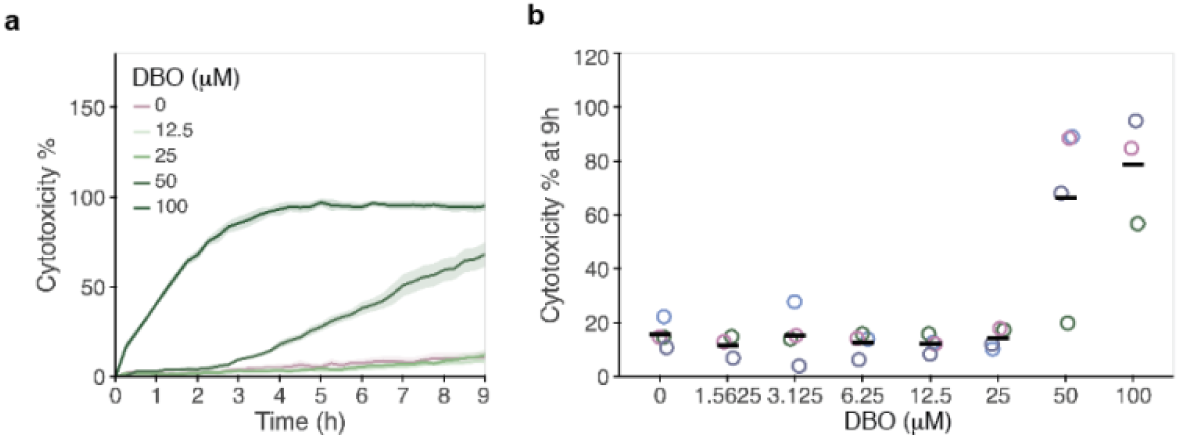
Cytotoxicity of dalberoside (DBO) in differentiated THP-1 cells. **a,** Representative cytotoxicity time courses following treatment with the indicated DBO concentrations. Lines and shading indicate the mean ± SEM of technical replicate wells. **b,** Cytotoxicity at 9 h after treatment. Colored open circles represent independent biological replicates and black horizontal lines indicate the mean; *n* = 3. Panel **a** shows one representative experiment from three biological replicates. Statistical analyses are described in Methods.

**Extended Data Fig. 6.**
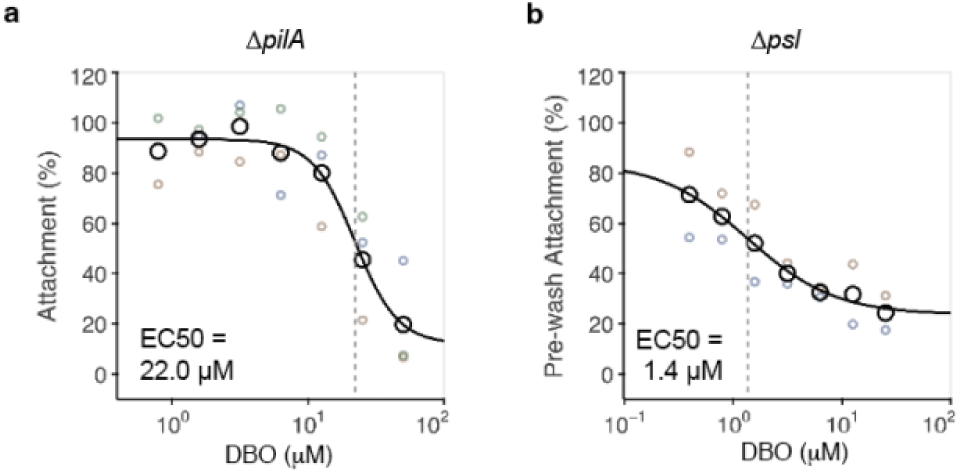
Dalberoside (DBO) inhibits attachment in attachment-defective *P. aeruginosa* backgrounds. **a,b,** Concentration–response analysis in Δ*pilA* using the post-wash assay (**a**) and Δ*psl* using the pre-wash assay (**b**). Cells were treated with the indicated DBO concentrations for 30 min. Attachment was quantified as total segmented cell area per image and normalized to the DMSO control within each biological replicate. Colored open circles indicate biological-replicate means, black open circles indicate means across replicates, black curves indicate four-parameter log-logistic fits to biological-replicate means and gray dashed lines indicate fitted EC50 values; *n* = 3 independent biological replicates.

**Extended Data Fig. 7.**
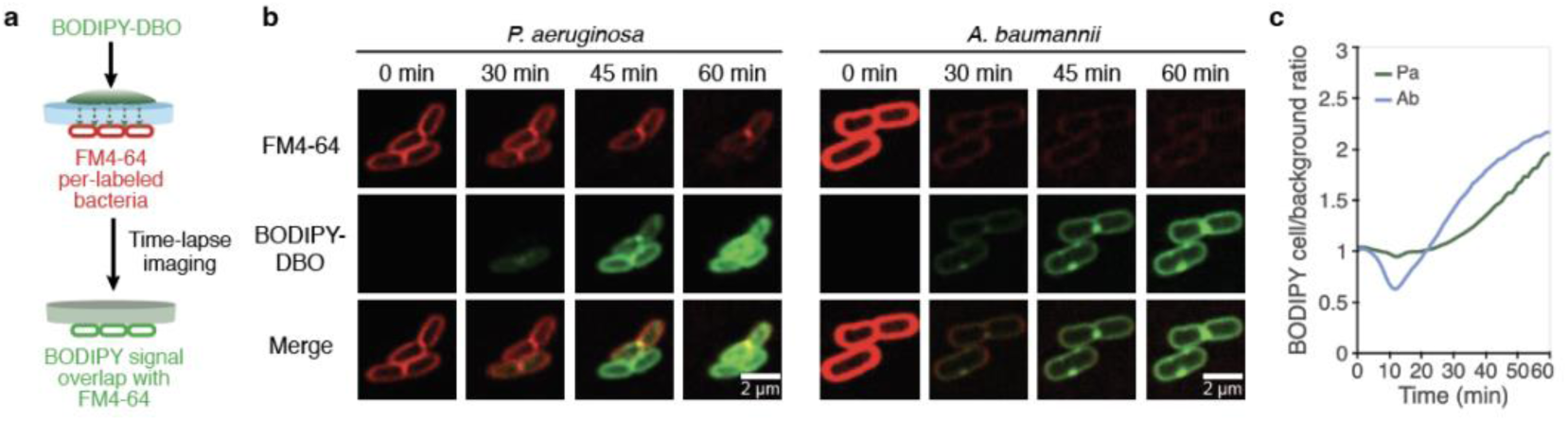
Time-lapse analysis of BODIPY–DBO association with bacterial cells. **a,** Schematic of the agarose pad–based labeling assay. FM4-64-stained bacteria were immobilized under an agarose pad and exposed to BODIPY–DBO for time-lapse imaging. **b,** Representative images of *P. aeruginosa* and *A. baumannii* at 0, 30, 45 and 60 min after BODIPY–DBO addition. FM4-64 and BODIPY–DBO are shown in red and green, respectively. **c,** Representative time course of cell-associated BODIPY–DBO fluorescence over 60 min, expressed relative to the mean field background outside buffered cell-associated regions.

**Extended Data Fig. 8.**
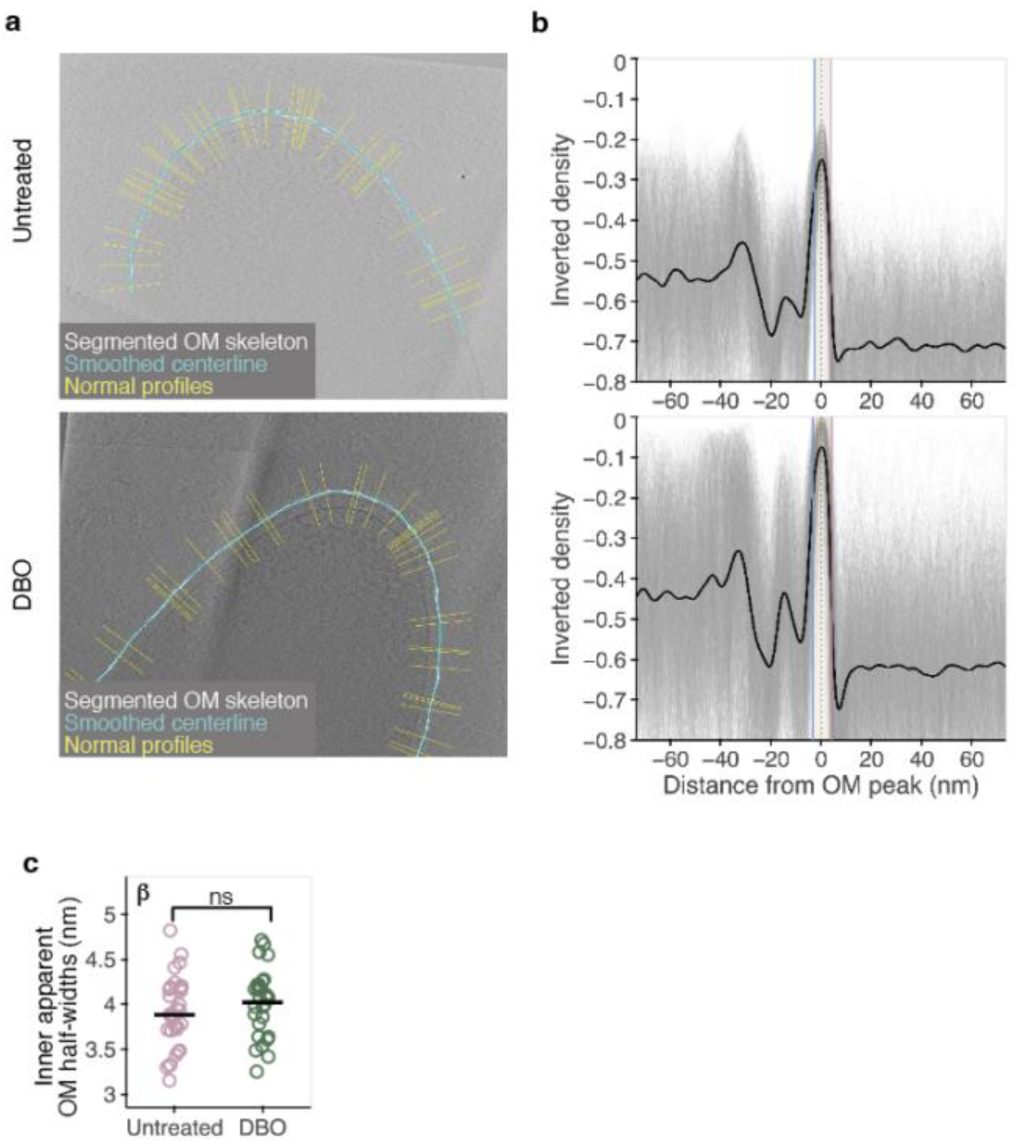
Representative cryo-ET outer membrane segmentation and density-profile analysis. **a,** Representative analysis of *P. aeruginosa* treated with DMSO or 25 µM DBO. White points indicate the segmented outer membrane skeleton, cyan lines the smoothed centerline and yellow lines the normals used for density extraction. **b,** Representative aligned inverted-density profiles relative to the outer membrane peak. Gray lines indicate individual profiles and the black line their mean. Outer-facing and inner-facing edges were defined by local maxima in the absolute gradient of the smoothed mean profile and are shown in red and blue, respectively. **c,** Inner-facing half-width β in DMSO- and DBO-treated cells. DMSO, *n* = 31 cells; DBO, *n* = 30 cells, sampled from one independently prepared specimen per condition. Statistical analyses are described in Methods.

